# Cryo-EM structures reveal the PP2A-B55α and Eya3 interaction that can be disrupted by a peptide inhibitor

**DOI:** 10.1101/2025.02.04.636346

**Authors:** Shasha Shi, Xueni Li, Christopher Alderman, Wei Huang, Lars Wick, North Foulon, John Rossi, Wenxin Hu, Shouqing Cui, Hongjin Zheng, Derek J. Taylor, Heide L. Ford, Rui Zhao

## Abstract

We have previously shown that Eya3 recruits PP2A-B55α to dephosphorylate pT58 on Myc, increasing Myc stability and enhancing primary tumor growth of triple-negative breast cancer (TNBC). However, the molecular details of how Eya3 recruits PP2A-B55α remain unclear. Here we determined the cryo-EM structures of PP2A-B55α bound with Eya3, with an inhibitory peptide B55i, and in its unbound state. These studies demonstrate that Eya3 binds B55α through an extended peptide in the NTD of Eya3. The Eya3 peptide and other PP2A-B55α substrates and protein/peptide inhibitors including B55i bind to a similar area on the B55α surface but the molecular details of the binding differ. We further demonstrated that the B55i peptide inhibits the B55α and Eya3 interaction in vitro. B55i peptide expressed on a plasmid increases pT58 and decreases Myc protein level in TNBC cells, suggesting the potential of B55i or similar peptides as therapies for TNBC.

## Introduction

The Eya family of proteins consists of four family members (Eya1-4) (1) that were originally identified as transcriptional co-activators of Six1 (2, 3). The Six1 and Eya transcriptional complex is critical for the development of eyes and several other organ systems in mammals (4-12). Six1 and Eya are typically downregulated after development is complete but are re-expressed in cancers (5, 8, 13-20). As the activator of the Six1/Eya bipartite transcription factor complex, Eya mediates critical functions in cancer by promoting proliferation, angiogenesis, plasticity, immune evasion, invasion, metastasis, and epithelial-mesenchymal transition (EMT) (4, 5, 14, 21-24). The Eya protein also has transcription-independent roles that are orchestrated through its C-Terminal Domain (CTD), which contains intrinsic tyrosine phosphatase activity (8, 25, 26), as well as its N-Terminal Domain (NTD), which contains S/T phosphatase activity through its interaction with PP2A-B55α (8, 27-29). Both the tyrosine phosphatase and S/T phosphatase activities contribute to oncogenic processes of Eya proteins that can be independent of Six1. For example, the tyrosine phosphatase mediates transcription (30), functions in DNA damage response (31-33), regulates the cytoskeleton (23, 34), promotes mitotic progression and centrosome maturation (35, 36), induces angiogenesis (21, 37, 38), regulates ERβ signaling (30), promotes Myc stabilization (8, 39), increases tumor metastasis (23, 40), and maintains progenitor cell populations (35). Similarly, the S/T phosphatase upregulates PD-L1 (41), stabilizes Myc (4, 27, 42, 43), drives metastasis (44), allows replication fork progression (44), and regulates innate immune signaling (45). Since Six1 and Eya are typically downregulated in healthy tissues after development is complete, they represent ideal targets for new cancer therapies (13, 14).

Eya proteins have recently been shown to play an important role in Myc stabilization through Myc pT58 dephosphorylation (46). Myc is one of the most prolific proto-oncogenes (47-53) due to its ability to drive several cancer hallmarks (47, 54, 55). Thus, under normal conditions, Myc is tightly regulated to avoid its accumulation and the induction of cancer. Myc can be regulated at the level of transcription through transcription factors, DNA structures, and promoter elements (56-60). It can also be regulated at the level of translation or post-translation through microRNAs (61-65), phosphorylation, prolyl isomerization, and ubiquitination to change the Myc protein half-life (66-71). Dysregulation of any of these Myc regulatory networks often leads to oncogenesis (66, 70-76). While it was previously believed that Eya dephosphorylates Myc pT58 directly (42), we have shown that Eya3-mediated pT58 dephosphorylation and stabilization of Myc involves the recruitment of PP2A-B55α (27).

PP2A is a major Ser/Thr phosphatase in the cell (77, 78) and is generally assembled as a trimeric enzyme composed of a 65 kD structural subunit (A), a 36 kD catalytic subunit (C), and a regulatory subunit (B) of varying sizes. The A and C subunits interact to form the dimeric core enzyme, which only gains its full activity, subcellular localization, and substrate specificity after interacting with one of the many regulatory B subunits to form the trimeric holoenzyme. B subunits are extraordinarily complex with at least 26 B subunits identified in humans that can be divided into 4 subfamilies, B (B55/PR55), B’ (B56/PR61), B” (B72/PR72), and B”’ (PR93/PR110). In contrast to kinases, PP2A does not use residues around the active site of its catalytic domain to recognize specific sequences around the phosphosite to achieve substrate specificity (79-81). Instead, PP2A depends on its B subunit to recognize and bind to specific sequence motifs that are often distal from the phosphosite but help to position it in the active site of the catalytic C subunit of the PP2A heterotrimer (79-81).

We have shown that Eya proteins directly interact with the PP2A-B55α but not B56 subunit. Indeed, a PP2A-B55α binding mutant, H79A^Eya3^, impeded Myc stabilization and triple-negative breast cancer (TNBC) tumor metastasis (27). We further showed that Eya3 bridges PP2A-B55α and Myc, using its intrinsically disordered NTD between residues 53-90 to recruit PP2A-B55α, and uses its CTD to bind the NTD of Myc (46). Moreover, all Eya family members can interact with PP2A-B55α through analogous disordered regions in their NTDs (27, 46). We also showed Eya3 binds to an area of PP2A-B55α that partially overlaps with its substrate-binding surface (46).

These observations suggest that disrupting the PP2A-B55α and Eya3 interaction may be a useful approach to inhibit Myc stabilization in tumors that over-express Eya3. Kruse and colleagues have used protein engineering to design peptides that bind the proposed substrate binding surface of B55 and identified one of the peptides, B55i, as a strong binder of PP2A-B55α that inhibits its interaction with substrates (82). Here, we have determined the cryo-EM structures of Eya3 or B55i bound to PP2A-B55α. These structures revealed the molecular details of the Eya3 and B55α interaction and showed the B55i binding site on B55α overlaps with that of Eya3. We showed B55i can disrupt the Eya3 and PP2A-B55α interaction in vitro, and B55i expressed on a plasmid can increase pT58 and reduce Myc protein level in TNBC cells, illustrating its potential as a cancer therapy.

## Results

### Structure of the PP2A-B55 and Eya3 complex

We expressed and purified Eya3 protein from *E. coli* and the His-B55α subunit from insect cell culture (**Supplementary Fig. 1**). In the remainder of the paper, we will use B55 to represent B55α for simplicity and PP2A-B55 to represent the holoenzyme containing the A, B55, and C subunits. We labeled His-B55 with the RED-tris-NTA dye and carried out microscale thermophoresis (MST) experiments with increasing concentrations of Eya3. This experiment demonstrated that Eya3 binds to B55α with a Kd of 1.04 μM (**Fig. 1A**).

**Fig. 1.**
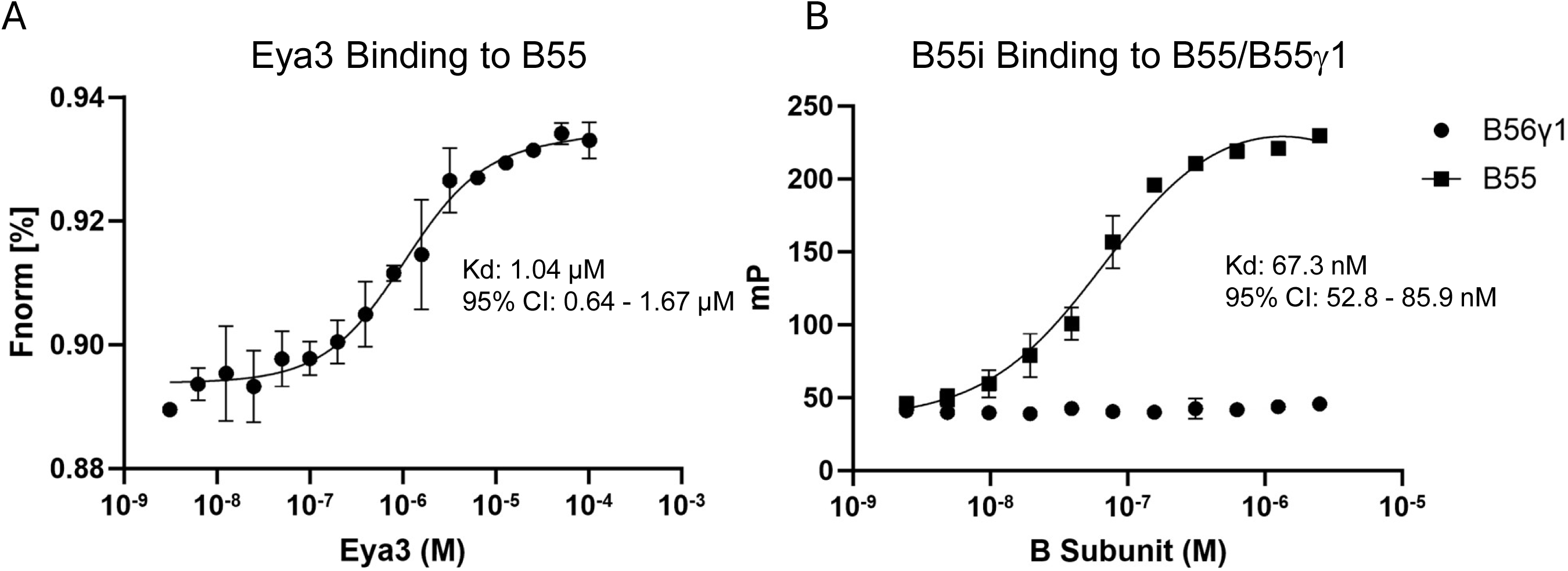
Purified Eya3 and B55i bind PP2A-B55. A) MST experiment showing the binding affinity between Eya3 and B55. Error bars represent the standard deviation. The dataset represents three independent biological replicates. Kd and 95% confidence interval (CI) are shown. B) Fluorescence polarization experiment showing the binding affinity between B55i and B55 or B56γ1. Error bars represent the standard deviation. The dataset represents three independent biological replicates.

To determine the PP2A-B55 + Eya3 structure, we over-expressed Flag-B55 in HEK293 FreeStyle cells. We pulled down Flag-B55 using anti-Flag resin which also brought down the endogenous A and C subunits and eluted the complex using Flag peptide (**Supplementary Fig. 2A**). We then incubated the PP2A-B55 complex containing all three subunits with full-length Eya3 purified from HEK293 FreeStyle cells (**Supplementary Fig. 2B**), evaluated the complex on negative stain grids (**Supplementary Fig. 2C**), prepared cryo-grids (**Supplementary Fig. 2D, E**). We determined the cryo-EM structure of the PP2A-B55 + Eya3 complex to an overall resolution of 3.7 Å (**Supplementary Fig. 3A-C, Supplementary Table 1**).

The structure of PP2A-B55 + Eya3 shows the typical PP2A architecture with its horseshoe-shaped structural A subunit interacting with the more globular regulatory B subunit (which is a WD domain made of a 7-blade β-propeller) and catalytic C subunit (**Fig. 2A**). The C-terminal tail of C subunit (residues 294 to 309) is clearly visible (**Fig. 2A**). The middle portion of the C-terminal tail (residues 298 to 303) interacts extensively with residues on β3D of blade 3 in B55, and the end of the C-terminal tail (residues 304-309) extends to the interface between the A and B55 subunits and interacts with both subunits (**Fig. 2B**). The conformation and interactions of the C-terminal tail are similar to those in the structures of PP2A-B55 + ARPP19 and PP2A-B55 + FAM122A (83) but different from the crystal structure of PP2A-B55 + microcystin-LR (MCLR) in which the C-terminal tail of C subunit is not visible (79).

**Fig. 2.**
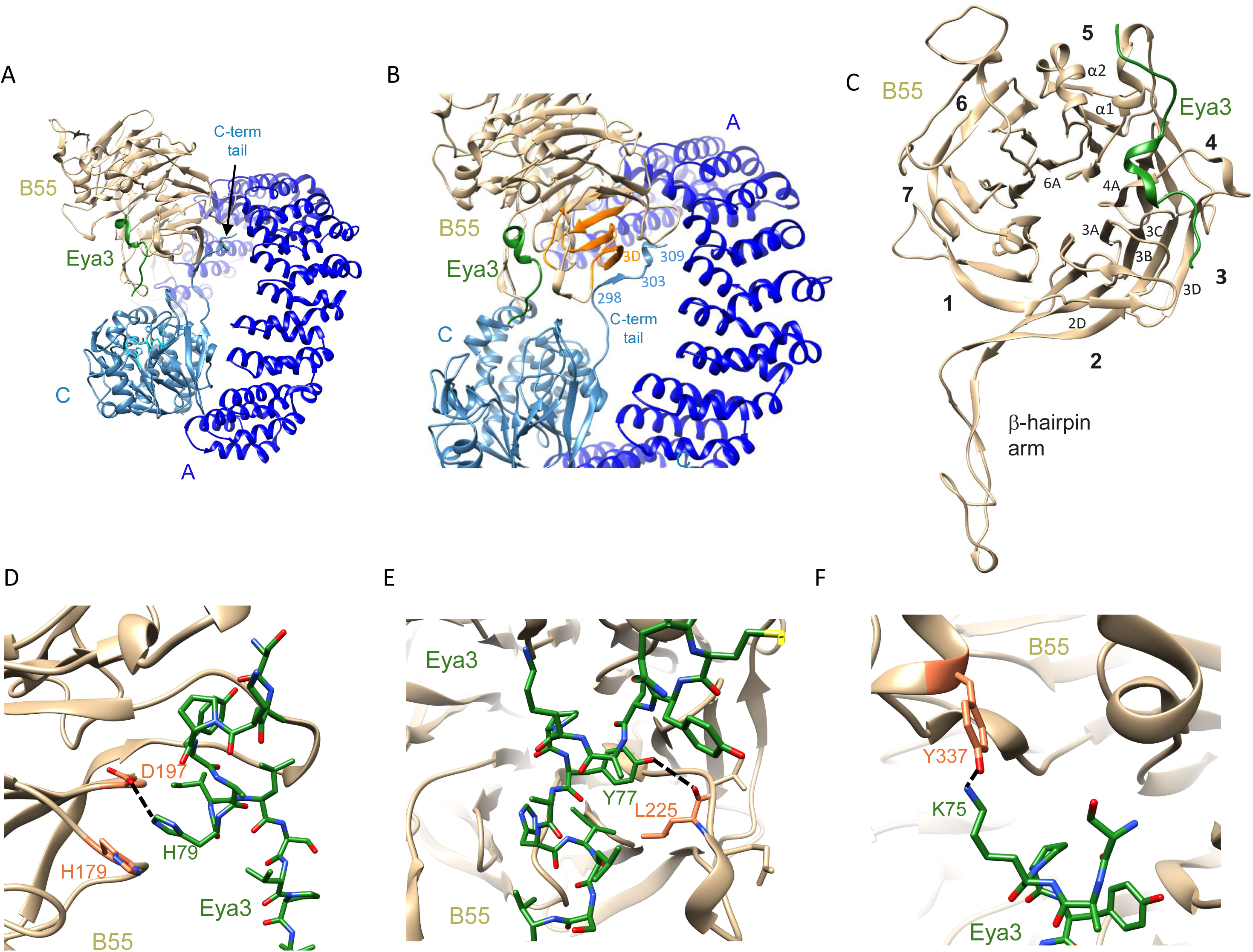
Cryo-EM structure of the PP2A-B55 + Eya3 complex. A) The overall structure of the PP2A-B55 + Eya3 complex. B) A zoom in view of panel A showing the middle portion of the C-terminal tail (residues 298-303) of C subunit interacts with the β3D strand of blade 3 (orange) in B55. It also shows that the end of the C-terminal tail (residues 304-309) inserts into the A and B55 subunit interface and interacts with both subunits. C) The structure of the B55 subunit with Eya3 bound. The blades of the β-propeller are labeled with numbers. The β-sheets and α-helices involved in Eya3 interaction are also labeled. D) Residue H79^Eya3^ forms a hydrogen bond with D197^B55^ and pi-pi or pi-cation interaction with H179^B55^. E) Residue Y77^Eya3^ forms a hydrogen bond with the main chain carbonyl oxygen of L225^B55^. F) Residue K75^Eya3^ forms a hydrogen bond with Y337^B55^.

Despite the presence of full length Eya3 in the complex, only density for a stretch of extended peptide is visible in the structure (**Fig. 2A)**. Using the information from our previous biochemical studies that a peptide between residues 53 and 90 in Eya3 binds to B55 (46), we can model residues 73-79 into this stretch of density (**Supplementary Fig. 3D**). Weaker density up and downstream of these residues can be modeled for the main chain of Eya3 residues 65-72 and 80-86, but side chain conformations for these residues are less defined.

The modeled Eya3 peptide (residues 65-86) interacts mostly with residues on blades 3 and 4 of the β-propeller on the B55 surface (including residues on the β2D and β3A loop, β3B and β3C loop, β3D and β4A loop) and with residues on the α1 and α2 helix positioned above blade 5, as well as the α2 and β6A loop (**Fig. 2C**). In particular, residue H79^Eya3^ forms pi-pi or pi-cation stacking with H179^B55^ and a hydrogen-bond with D197^B55^ (**Fig. 2D**). The H79A^Eya3^ mutant has been shown to dramatically reduce the interaction between Eya3 and B55 (28, 46). Furthermore, Y77^Eya3^ forms a hydrogen bond with the main chain carbonyl of L225^B55^ and K75^Eya3^ forms another hydrogen bond with Y337^B55^ (**Fig. 2E and F**). The Y77A^Eya3^ mutant has been shown to abolish the apparent Ser/Thr phosphatase activity of Eya3 (28), which is likely caused by the disruption of the Eya3 and PP2A-B55 interaction. While we were working on the structure of the PP2A-B55 + Eya3 structure, Padi and colleagues published the cryo-EM structure of PP2A-B55 in complex with an Eya3 peptide (residues 62-108) on bioRxiv (84). Although the pdb file has not yet been released and we cannot perform a detailed comparison with our structure, the description in the bioRxiv paper indicates that the two structures are similar.

### Structure of the PP2A-B55 and B55i complex

Kruse and colleagues have computationally engineered a B55i peptide that binds to B55 and inhibits its interaction with several B55 substrates (82). In our hands, the synthesized B55i peptide binds B55 purified from insect cells with a Kd of 67.3 nM but does not bind B56γ1 in fluorescence polarization experiments (**Fig. 1B**).

We next incubated the PP2A-B55 complex purified from HEK293 FreeStyle cells with the B55i peptide and determined its cryo-EM structure to an overall resolution of 3.5 Å (**Supplementary Fig. 4A-F, Supplementary Table 1**). The C-terminal tail of the C subunit is clearly visible, and its conformation and interactions are similar to that in the PP2A-B55 + Eya3 complex. The structure also shows that B55i forms a helix-turn-helix structure, as predicted by Kruse and colleagues (82). Density for the C-terminal helix of B55i (residues 21-32) is well defined (**Supplementary Fig. 4G**), but the density for the N-terminal helix is much weaker. Although the main chain of the N-terminal helix can be modeled, the side chain conformations are less defined (other than K10) and the density for the connecting loop between the N and C-terminal helix is even weaker.

B55i binds to a similar surface as the Eya3 peptide, mostly contacting blades 3 and 4 of the β-propeller and the α1 and α2 helices positioned above blade 5 on the B55 surface (**Fig. 3A, B**). In particular, K10^B55i^ forms a salt bridge with D340^B55^ and a hydrogen bond with the carbonyl oxygen of Y337^B55^ (**Fig. 3C**). Residue K23^B55i^ forms a hydrogen bond with D340^B55^ and its aliphatic chain forms hydrophobic interactions with F343^B55^ (**Fig. 3D**). Residue R38^B55i^ forms a hydrogen bond with E223^B55^ (**Fig. 3E**). The helix-loop-helix conformation of B55i and its binding surface on B55 are also very similar to that of the protein inhibitor IER5 (85).

**Fig. 3.**
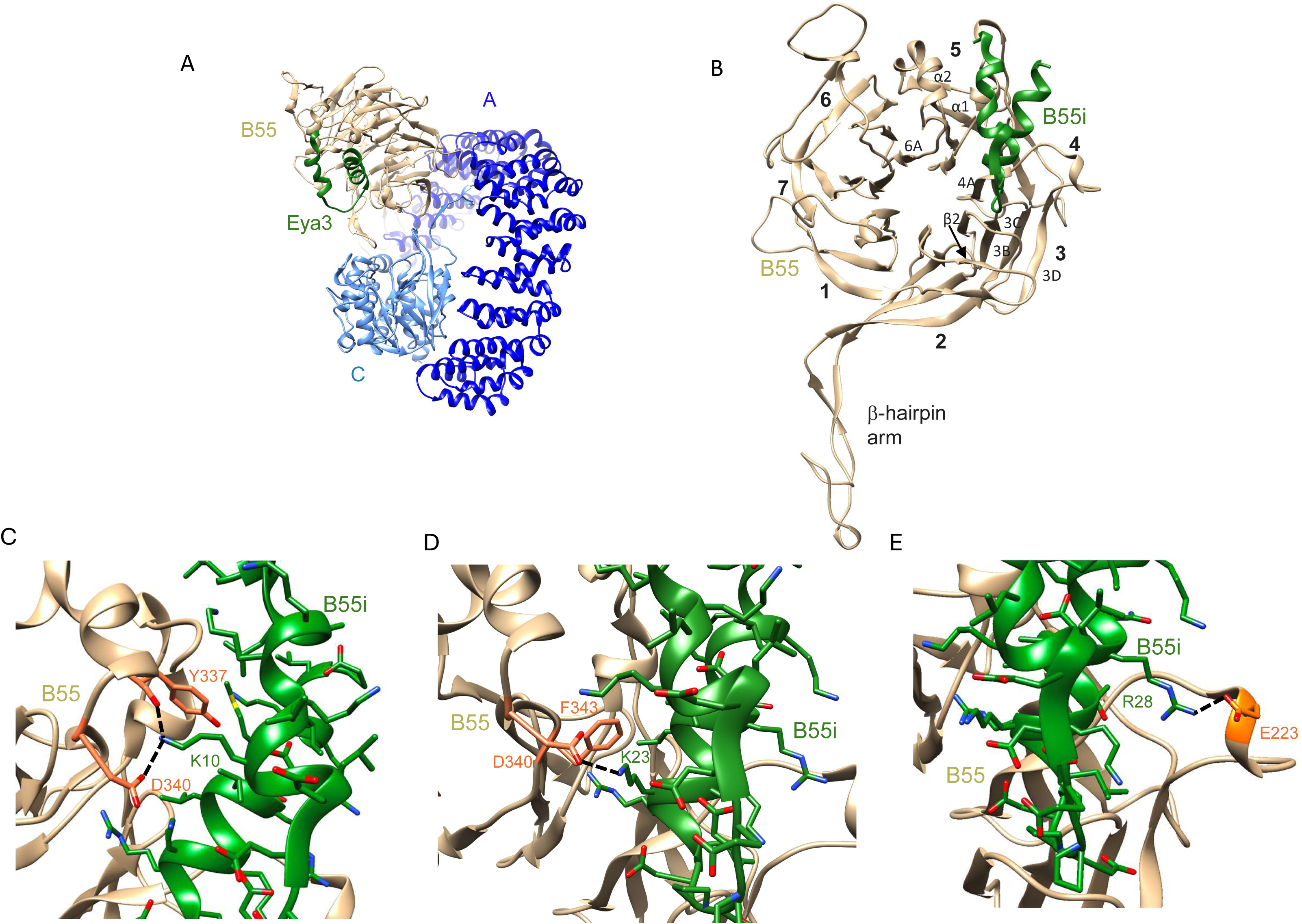
Cryo-EM structure of the PP2A-B55 + B55i complex. A) The overall structure of the PP2A-B55 + B55i complex. B) The structure of the B55 subunit with B55i bound. The blades of the β-propeller are labeled with numbers. The β-sheets and α-helices involved in B55i interaction are also labeled. C) Residue K10^B55i^ forms a salt bridge with D340^B55^ and a hydrogen bond with the carbonyl oxygen of Y337^B55^. D) Residue K23^B55i^ forms a hydrogen bond with D340^B55^ and its aliphatic chain forms hydrophobic interaction with F343. E) Residue R28^B55i^ forms a hydrogen bond with E223^B55^.

### Structure of the apo PP2A-B55 complex

There are currently several solved PP2A-B55 structures, including the crystal structure of PP2A-B55 + MCLR (a cyclic peptide inhibitor that binds to the active site in the C subunit (79)), the cryo-EM structures of PP2A-B55 + phosphorylated ARPP19 (a protein inhibitor that binds to both the B and C subunits (83)), PP2A-B55 + NTD of FAM122A (a protein inhibitor that binds to both the B and C subunits (83)), PP2A-B55 + NTD of IER5 (a protein inhibitor that only contacts the B subunit (85)), PP2A-B55 + p107 (a substrate that mainly binds to the B subunit with its C-terminal end extending towards, but not contacting, the C subunit (84)), and PP2A-B55 + Eya3 peptide (residues 62-108 (84)). However, there is no reported apo PP2A-B55 structure. To fill this void, we determined the cryo-EM structure of the PP2A-B55 complex on its own to an overall resolution of 3.2 Å (**Supplementary Fig. 5A-F, Supplementary Table 1**).

The first notable feature of the apo PP2A-B55 structure is that the middle portion (residues 294 to 303) of the C-terminal tail of the C-subunit do not have visible density, but the last few residues (304 to 309) of the C-terminal tail have well defined density with the same conformation as in the Eya3 or B55i bound PP2A-B55 structure and are nestled between the A and B subunits (**Supplementary Fig. 5G**). This is different from the crystal structure of PP2A-B55 bound to MCLR and all the other cryo-EM structures of PP2A-B55 bound to a substrate/inhibitor/recruiter (**Fig. 4A**). The crystal structure of PP2A-B55 bound to MCLR lacks density for the entire C-terminal tail of the C subunit. All the other cryo-EM structures of PP2A-B55 with a binder (substrate/inhibitor/recruiter) have well defined density for the entire C-terminal tail of the C subunit **(Supplementary Fig. 5G)**.

**Fig. 4.**
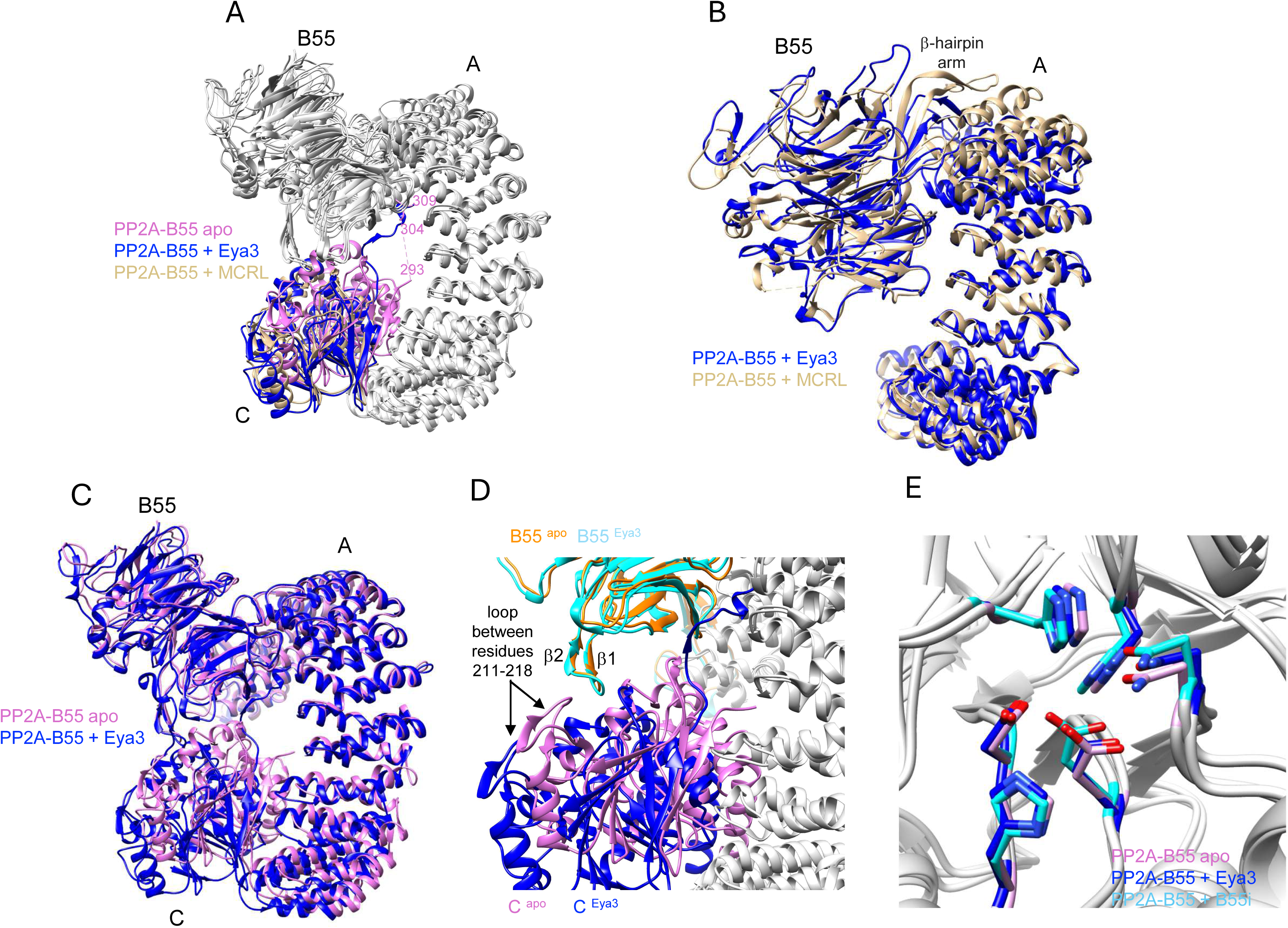
Cryo-EM structure of the apo PP2A-B55 complex. A) The structures of the PP2A-B55 apo, PP2A-B55 + Eya3, and PP2A-B55 + MCRL superimposed on their A subunit. Only the C-subunit is colored for clarity. The C-terminal tail (residues 294-309) of the C-subunit is present in the Eya3 structure, absent in the MCRL bound structure, and partially (residues 304-309) present in the apo structure. B) The PP2A-B55 + Eya3 and PP2A-B55 + MCRL structures superimposed on their A subunit. Only the B and C subunits are shown for clarity. The N-terminal region of A subunit is more contracted and the β-hairpin arm of B55 is closer to the A subunit in the Eya3 structure than the MCRL structure. C) The PP2A-B55 + Eya3 and PP2A-B55 apo structures superimposed on their A subunit. The C-terminal region of A subunit is more contracted, and the C subunit is closer to A and B55 in the apo structure than in the Eya3 structure. D) A zoom in view of the interface between the B55 and C subunit in the apo and PP2A-B55 + Eya3 structure showing that the loop between residues 211-218 in the C subunit and the β1-β2 hairpin of B55 are close and interact with each other in the apo structure but are far apart in the Eya3 structure. A subunit in both structures are shown in grey for clarify. E) Active site residues of apo PP2A-B55, PP2A-B55 + Eya3, and PP2A-B55 + B55i with the C subunit superimposed show no significant differences.

The second notable feature of the apo PP2A-B55 structure is the different curvature of the horseshoe-shaped A subunit and spatial organization of its three subunits compared to the other PP2A-B55 structures. It was observed previously that the binding of a protein inhibitor (ARPP19, FAM122A, IER5), substrate (p107), or recruiter (Eya3) make the A subunit more contracted in its N-terminal region compared to the crystal structure of PP2A-B55 bound to MCLR (79, 83-85). Our Eya3 and B55i bound structures as well as the apo structure follow the same trend with their N-terminal region of A subunit being more contracted compared to the crystal structure of PP2A-B55 bound to MCLR (**Fig. 4B**, only the Eya3 and MCLR structural comparison is shown for clarity). However, the C-terminal region of the A subunit in the apo structure is also more contracted, and its C subunit is substantially closer to the B55 and A subunits than in any other PP2A-B55 structures, making the apo PP2A-B55 the most compact structure among all PP2A-B55 structures determined (**Fig. 4C**, only the Eya3 and apo structural comparison is shown for clarity). In this compact apo structure, the loop between residues 211-218 in C subunit is brought closer to and interacts with the β1-β2 hairpin of B55 (**Fig. 4D**).

### PP2A-B55 binders inhibit its intrinsic phosphatase activity

ARPP19 and FAM122A have been shown to inhibit the phosphatase activity of PP2A-B55 in vitro using a small molecule substrate DiFMUP (83). It is not surprising that ARPP19 and FAM122A inhibit the PP2A-B55 phosphatase activity since these two proteins bind on the surface of both B55 and C subunits, blocking access to the active site on C subunit. Padi and colleagues have shown that p107 and Eya3 peptide also weakly inhibit the phosphatase activity of PP2A-B55 (IC_50_s for p107 and Eya3 are 245 +/- 27 nM and 1102 +/- 84 nM, respectively) (84), even though p107 and the Eya3 peptide only bind to the B55 subunit. In contrast, we have previously shown that Eya3 protein increases the phosphatase activity of PP2A-B55 in a malachite green assay using phosphorylated peptide substrates (46). We noted that the buffer condition we used in the malachite green assay (50 mM Tris pH 7.5, 50 mM NaCl, 5 mM MgCl_2_, and 0.1 mM EDTA) is substantially different from that used by Page and Peti (30 mM HEPES pH 7.0, 150 mM NaCl, 1 mM MnCl_2_, 1 mM DTT, 0.01% triton X-100, and 0.1 mg/ml BSA) (46, 84). Using the same phosphatase assay condition as Padi and colleagues, we showed that the Eya3 NTD indeed inhibits the phosphatase activity of PP2A-B55 using OMFP as a substrate (**Fig. 5A**). Since our B55i peptide contains a FITC label which interferes with the OMFP assay, we tested its effect in a malachite green assay using a modified phospho-Myc peptide as substrate and showed that B55i also inhibits the phosphatase activity of PP2A-B55 (**Fig. 5B**). These data show that all B55 binders studied so far weakly inhibit the intrinsic phosphatase activity of PP2A-B55, even if they only bind to the B55 subunit.

**Fig. 5.**
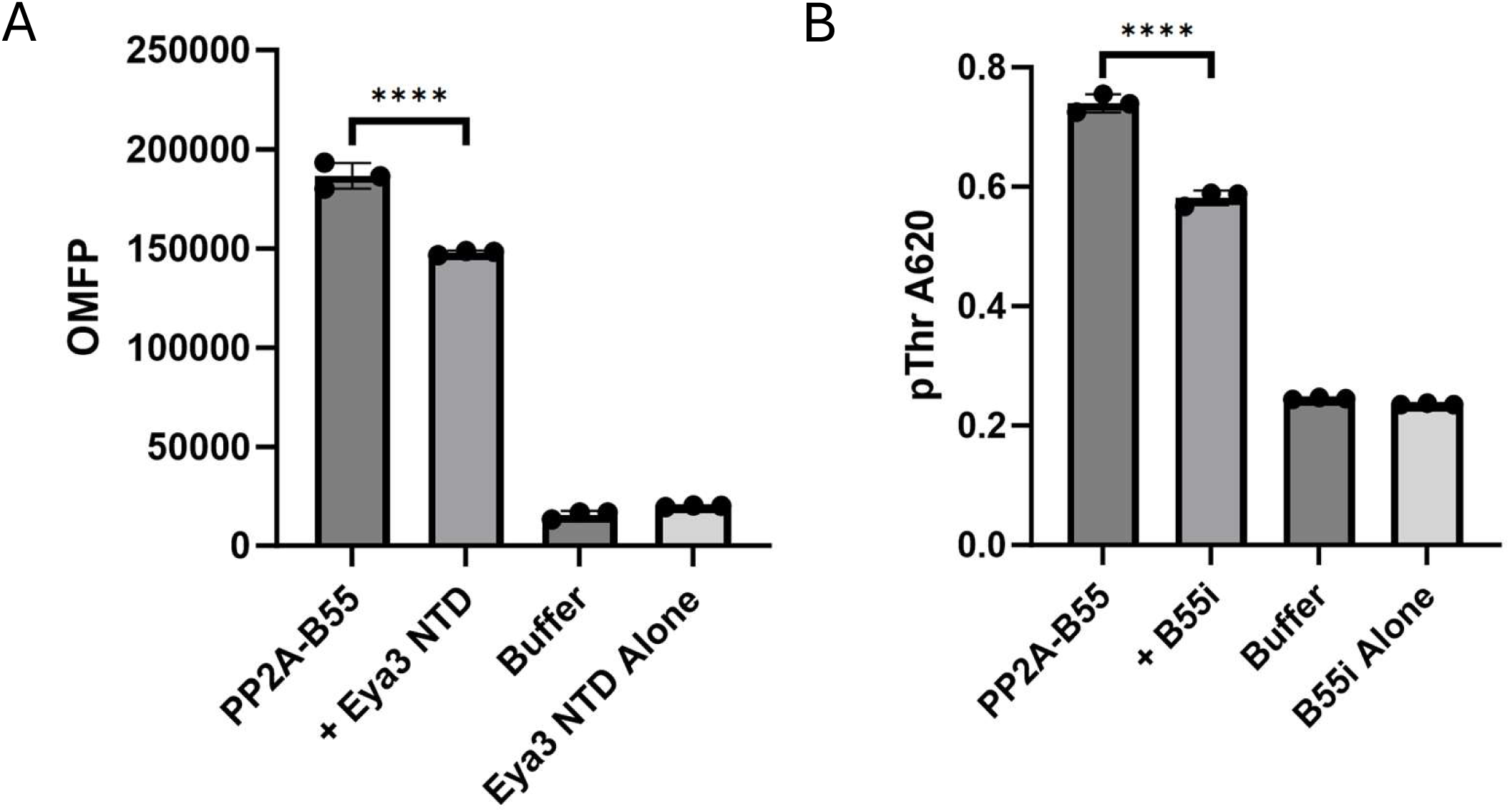
Eya3 NTD and B55i inhibit the phosphatase activity of PP2A-B55. A) The phosphatase activity of PP2A-B55 in the absence and presence of Eya3 NTD in a fluorescent phosphatase assay using OMFP as a substrate. B) The phosphatase activity of PP2A-B55 in the absence and presence of B55i peptide in a malachite green phosphatase assay using a modified Myc pT58 peptide. The dataset represents three independent biological replicates. Error bars represent the standard deviation. Statistical data analysis was done using Fmax tests followed by two-tailed, homoscedastic t-tests. ****P<0.001.

### PP2A-B55 substrate, inhibitors, and recruiter bind to similar areas on B55 with distinct specific interactions

We next compared the binding sites of Eya3, B55i, and two previously determined PP2A-B55 protein inhibitors (ARPP19 and FAM122A) whose structures have been deposited in the Protein Data Bank (83). The B subunits in all these structures are very similar (**Fig. 6A**). The four peptides share a core binding surface centered around blades 3-5 of the β-propeller on B55 (**Fig. 6B**). Both B55i and Eya3 bind exclusively on B55 with the C-terminus of Eya3 extending to the C subunit without making direct contact. This mode of binding seems to be the case for substrate p107 and another protein inhibitor, IER5, as well (84, 85). On the other hand, ARPP19, and FAM122A bind both the B55 and C subunits with their C-terminal halves blocking the active site surface on the C subunit. Despite binding a similar region on the surface of B55, the specific interactions are different. For example, the Eya3 peptide uses a Lys-x-Tyr-x-His stretch of residues as key contributors to its interaction with B55, while B55i uses Lys-X12-Lys-X4-Arg residues that are far apart from each other (not spatially overlapping with the Eya3 residues either) to interact with a different set of residues on B55.

**Fig. 6.**
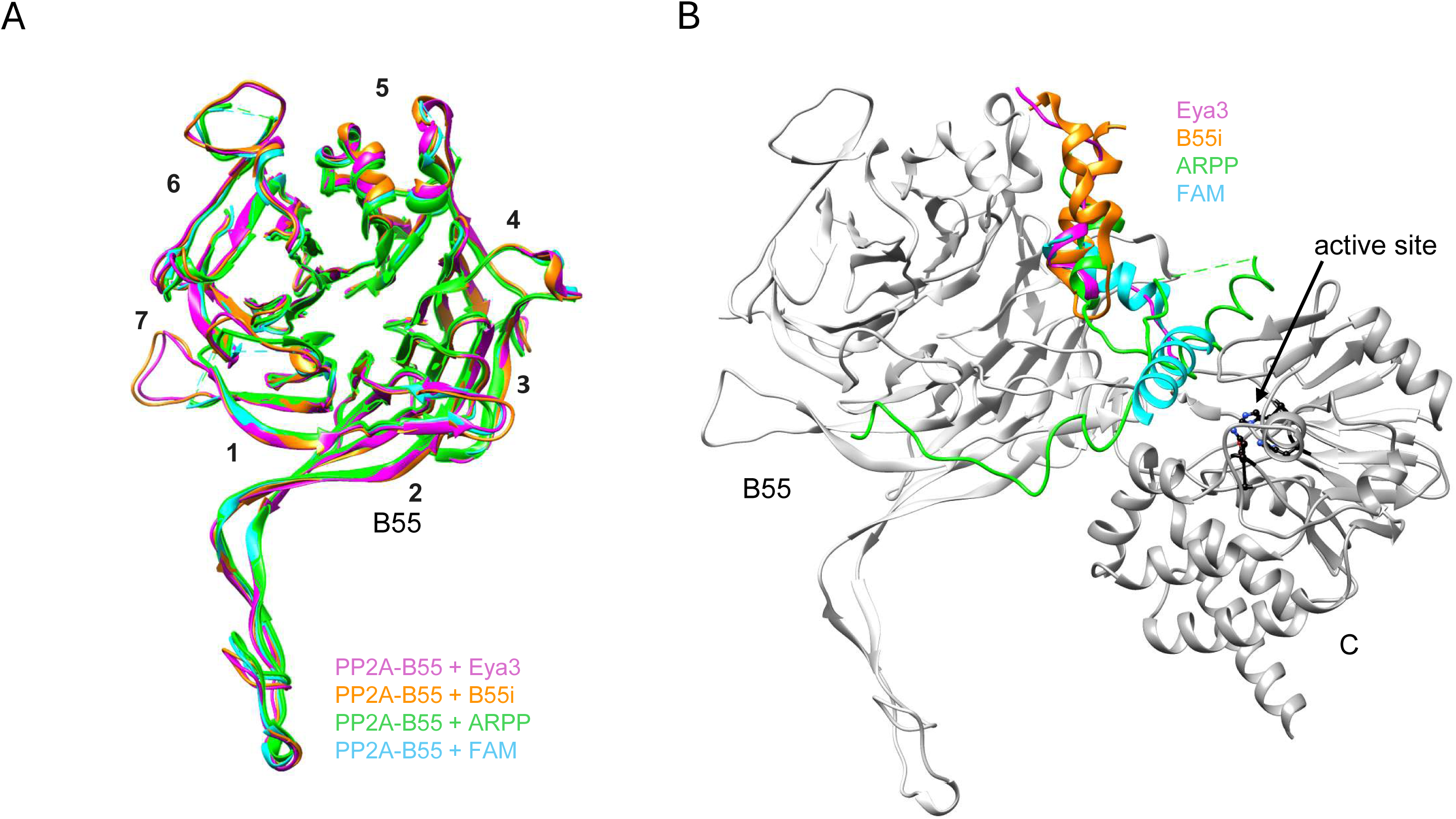
PP2A-B55 substrate/inhibitor/recruiter binds to similar areas on B55 with distinct specific interactions. A) Superimposition of the B55 subunit of PP2A-B55 + Eya3, + B55i, + ARPP19, or + FAM122A show the overall structure of the B55 subunit is very similar among the 4 complexes. B) Conformation of Eya3, B55i, ARPP19, and FAM122A on the surface of B55 with their B55 subunit superimposed (only the B55 and C subunit of Eya3 are shown in grey for simplicity).

### B55i inhibits B55 and Eya3 interaction

The structure of the PP2A-B55 and B55i complex suggests that B55i will inhibit the Eya3 and B55 interaction due to the overlapping binding sites of B55i and Eya3 on B55. Indeed, we showed that Eya3 and B55 form a complex in mass photometry (MP) experiments, and the addition of B55i disrupts this complex (**Fig. 7A**). We also showed that in the presence of B55i, Eya3 no longer binds to B55 in MST experiments (**Fig. 7B, C**). To evaluate the effect of B55i in cells, we expressed the B55i peptide as a GFP fusion on a plasmid in 66cl4 TNBC cells. We showed that GFP-B55i increased pT58 level while reduced Myc protein level (compared to GFP alone) in 66cl4 cells (**Fig. 7D**), consistent with the peptide disrupting the Eya3 and B55 interaction.

**Fig. 7.**
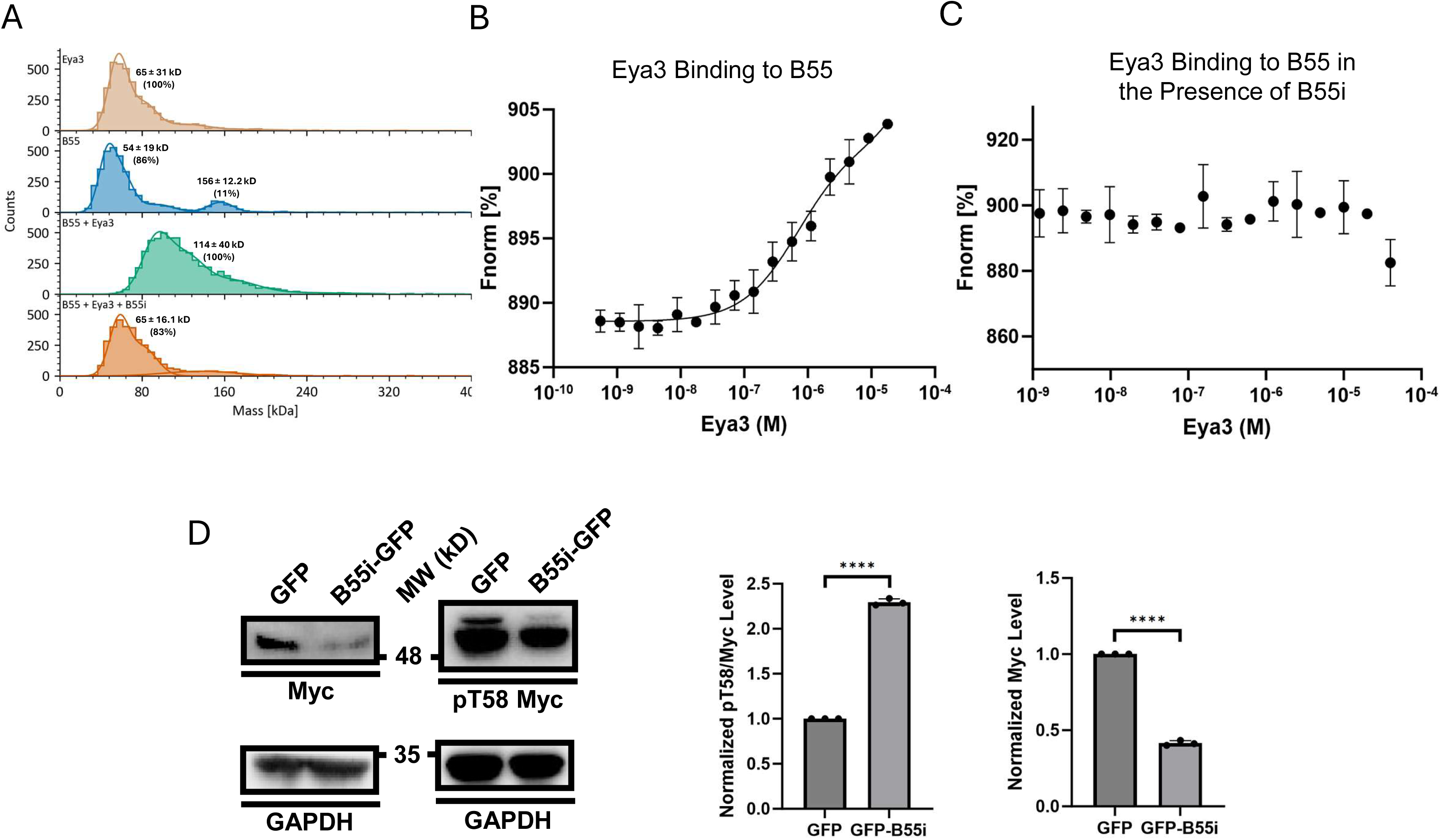
B55i inhibits the interaction between PP2A-B55 and Eya3. A) MP data showing the histograms of Flag-hEya3, His-B55, Flag-hEya3 + His-B55, and Flag-hEya3 + His-B55 + B55i, demonstrating that B55i disrupts Eya3 and B55 complex formation. B) MST binding affinity curve showing the interaction between mEya3 and His-B55α. C) B55i disruption of the mEya3 and His-B55α interaction on MST. D) B55i peptide expressed as a GFP-B55i fusion on a plasmid in 66cl4 TNBC cells led to induced pT58/Myc level and reduced Myc protein level as demonstrated by Western blot of pT58 and Myc protein level in whole cell lysate. The dataset in all panels represents three independent biological replicates. Error bars represent the standard deviation. Statistical data analysis was done using Fmax tests followed by two-tailed, homoscedastic t-tests. *P<0.05, **P<0.01, ****P<0.0001.

## Discussion

We have previously shown that Eya3 uses a peptide in its disordered NTD to interact with B55 and recruit PP2A-B55 to Myc, thereby dephosphorylating pT58 and stabilizing Myc (46). However, the molecular details of how Eya3 interacts with B55 remained unknown. Here we have determined the cryo-EM structure of the PP2A-B55 and full-length Eya3 complex, showing that residues 65-86 of Eya3 interact with mainly blades 3 and 4 as well as α1 and α2 positioned above blade 5 of the β-propeller on B55, similar to a recent cryo-EM structure of PP2A-B55 in complex with the Eya3 peptide (residues 62-108) reported by Padi and colleagues on bioRxiv (84), and consistent with our and others’ biochemical observations (28, 46, 82). In addition, we also determined the cryo-EM structure of PP2A-B55 in complex with a computationally engineered B55i peptide (82) and the apo PP2A-B55 complex.

Comparison of the apo and other PP2A-B55 structures (which are all bound structures containing a substrate, inhibitor, or recruiter) revealed interesting differences of the methylated C-terminal tail of the C subunit. The C-terminal L309 in the C subunit is reversibly methylated, and its methylation is required for PP2A assembly in cells (86-93) but not in vitro (79, 94, 95). In the crystal structure of PP2A-B55 + MCLR, the methylated C-terminal tail of PP2A-C subunit is disordered and invisible, while it is visible in all other cryo-EM structures of PP2A-B55 bound to a substrate, inhibitor, or recruiter (83-85, 94) and it extends to the A and B subunit interface. In the apo PP2A-B55 structure, the C-terminal tail is partially visible (the middle portion of the tail including residues 294-303 are not visible while the end of the tail including residues 304-309 are). The very C-terminal end of the tail (residues 304-309) is present and nestled between the A and B subunit in all PP2A-B55 structures (apo or bound) other than the MCLR-bound structure. In the latter, MCLR binds to the C subunit and disrupts its interaction with the β1-β2 hairpin of B55, which potentially leads to the slight changes between the B55 and A subunit interface (79). For example, the β-hairpin arm in B55 is much closer to the A subunit in the MCLR structure than in the Eya3 structure (**Fig. 4B**). These changes at the A and B55 interface may have prevented the very C-terminal end of the tail from binding to the A and B55 interface. We observed that the middle portion of the C-terminal tail of the C subunit interacts extensively with residues on blade 3 of B55 (**Fig. 2B**). Additionally, B55 substrate/inhibitor/recruiter binds to the same blade, though not to the exact β strand that the C-terminal tail interacts with (**Fig. 2B**). It is possible that B55 binders allosterically affect and enable the binding between the C-terminal tail and B55, which would explain why the middle portion of the C-terminal tail is visible in all substrate/inhibitor/recruiter bound PP2A-B55 structures but not in the PP2A-B55 apo structure.

The other notable difference between the apo and other PP2A-B55 structures is the curvature of the A subunit and the spatial organization of the three subunits. The N-terminal region of A subunit in all the cryo-EM structures is slightly more closed than that in the PP2A-B55 + MCLR crystal structure. We found that the apo structure is also more contracted in the C-terminal region of its A subunit and its C subunit is substantially closer to the A and B55 subunits compared to all the other PP2A-B55 structures (which are all bound structures). In the apo structure, the loop between residues 211-218 in the C subunit and the β1-β2 hairpin of B55 are close together and interact extensively, while the same interaction does not exist in the substrate/inhibitor/recruiter bound structures because the two components are too far apart (**Fig. 4D**). This interaction is likely responsible for making apo PP2A-B55 the most compact structure among PP2A-B55 complexes, which potentially led to the higher overall resolution of the apo structure compared to the PP2A-B55 + Eya3 and B55i structures. B55 binders potentially enable the binding of the middle portion of the C-terminal tail of C subunit to blade 3 of B55. This binding possibly hinders the C subunit from getting closer, leading to a more open and potentially more flexible structure. Indeed, local resolution analyses show that the apo structure has a more even resolution distribution (**Supplementary Fig. 5F**), while in the PP2A-B55 + Eya3 structures, the A and B55 subunit interface has the highest local resolution, while the C-terminal region of the A subunit and the C subunit have lower local resolutions (**Supplementary Fig. 3C**). These observations indicate that the A and B subunit interface is fairly rigid, while the C-terminal region of the A subunit and the C subunit have more flexibility. This suggests that the A subunit may serve as a stable anchor for B55 to capture the substrates through its substrate binding surface, while the C-terminal region of the A subunit and the C subunit are more dynamic, providing flexibility for the C subunit to access the phosphorylation site on the substrates that are likely at varied distances from the B55-binding site in different substrates.

These conformational changes caused by B55 binders may also explain the inhibition of intrinsic phosphatase activity towards small molecule substrates by B55i, Eya3, and substrate p107, even though they do not directly contact the C subunit, as shown here and by Padi and colleagues (84). These conformational changes do not change the active site substantially (**Fig. 4E**), but they may change the dynamics of the C subunit and the active site, which can affect enzymatic activity as observed in other systems (96). The weak inhibition of the PP2A-B55 activity by its B55 binders may have different consequences in the context of the natural function of these binders in the native environment of the cell. In the case of natural protein inhibitors such as ARPP19 and FAM122A, the inhibition of the intrinsic PP2A-B55 phosphatase activity through their binding to the B55 subunit augments their inhibitory function (which is mainly achieved through their binding to the C subunit and blocking of access to the active site). However, in the case of a substrate such as p107 and a recruiter such as Eya3, this weak inhibition is likely overcome by the dominant positive effect on dephosphorylation by their act of bringing the phosphosite close to the active site. For example, Eya3 clearly increases the dephosphorylation of pT58 on Myc and knocking down Eya3 reduces this dephosphorylation (27).

The mechanism of how PP2A-B55 recruits substrates has similarities and differences from PP2A-B56. The structure of B55 (a WD domain made of 7-blade β-propeller) is very different from B56 (which is composed of multiple HEAT repeats). B56 uses a pocket-like region to specifically bind a SLiM (Short Linear Motif) with the sequence of LxxIxE on substrate (97). On the other hand, B55 uses different residues to bind short peptide regions with diverse secondary structures (α-helix or extended peptide) and sequences. Of the six PP2A-B55 binders whose structures are determined now (ARPP19, FAM122A, IER5, p107, Eya3, and B55i), very few (p107 and FAM122A, and potentially Eya family members) share similar binding motifs, while the majority bind to B55 through very different sequence and interactions (79, 83-85). This diverse binding mechanism makes it difficult to predict the binding partners for B55 but also will likely enable B55 to bind many different regulators and recruiters. It will not be surprising to find other proteins like Eya3 that recruit different substrates that cannot be dephosphorylated by PP2A-B55 on their own. In both B55 and B56, the substrate binding site overlaps significantly with the binding site for regulators including protein inhibitors and recruiters (in the case of B56, the regulators and substrates bind to B56 using the same SLiM). This overlap suggests a major mechanism for protein inhibitors of both PP2A-B55 and B56 is to compete with substrate binding. In addition, PP2A-B55 inhibitors such as ARPP19 and FAM122A also inhibit PP2A-B55 activity by blocking access to the active site on the catalytic C subunit. The overlap between a recruiter (such as Eya3) binding surface on B55 and the substrate binding site also provides a unique advantage for recruiters such as Eya3 to achieve their function. It enables recruiters such as Eya3 to prevent B55 from binding other substrates while recruiting its preferred substrate (such as Myc) to be dephosphorylated by PP2A-B55. It is possible that the B subunits of the other two PP2A subfamilies (PP2A-PR72 and PP2A-PR93) also use the same or similar binding site for both substrates and regulators, making this competition model a general mechanism for regulating the PP2A families of phosphatases.

We demonstrated that the B55i peptide inhibits the Eya3 and B55 interaction and the B55i peptide expressed on a plasmid increases pT58 and reduces Myc protein level in TNBC cells (**Fig. 7**). This raised the interesting possibility of using B55i or a similar inhibitory peptide fused with cell penetrating peptide (CPP) as a potential therapy for TNBC. It is possible that the B55i peptide will interfere with PP2A-B55’s interaction with other cellular substrates. However, we have shown that B55 KD dramatically reduces TNBC metastasis without significant adverse effects in mice (27), suggesting that inhibiting the PP2A-B55 function will likely have a therapeutic window for breast cancer. Given that targeting Myc still remains a major unrealized goal for cancer therapy, the direct inhibition of the Eya3/PP2A interaction by CPP-B55i may be a significantly less toxic means to target Myc in TNBC and other tumor types that over-express Eya3.

## Experimental procedures

### Cloning

Flag-hB55, Flag-hEya3, GFP, and GFP-B55i were cloned into the pCAG vector; pGEX-6P-1 GST-mEya3 was cloned into pGEX-6p1; His-hB56γ1 was cloned into pGEX-4T1; and His-hB55 was cloned in pFASTbac, all using the NEBuilder HiFi DNA Assembly Cloning Kit (New England Biolabs). h and m are used to represent human and mouse proteins.

### Cell Culture

66cl4 murine TNBC and H1299 human non-small cell lung carcinoma cells were cultured using Dublecco’s Modified Eagle’s Media (DMEM) (Thermo Scientific) and RPMI-1640 (Thermo Scientific) supplemented with 10% FBS (Thermo Scientific) at 37 °C with 5% CO_2_. Cells were collected and validated as previously described (46).

### Protein Purification

GST-mEya3 was expressed in BL21 cells. Cells were lysed in lysis buffer containing 50 mM pH 7.5 Tris, 250 mM NaCl, 5% glycerol, 1 mM DTT, and 1X PIC. Lysis was done in ice with sonication using 4 x 45 second pulses with 2-minute off periods. NaCl was brought up to 0.5 M using 5 M NaCl and the lysate was centrifuged for 40 min at 18,000 G. Supernatant was gravity-flown over glutathione resin twice. Resin was washed with 100 mL of lysis bugger. GST was cleaved with PreScission protease and eluted into lysis buffer. Eluates were pooled and concentrated with Amicon Ultra Centrifugal filters. Concentrated protein was separated on a pre-equilibrated S200 column for size-exclusion chromatography. Peaks were combined, concentrated again, aliquoted, and stored at -80 °C.

FreeStyle 293-F cells were grown in SFMTransfx medium (Cytiva) at 37 °C and 8% CO2. For transient overexpression of B55 and h-Eya3 constructs, cells were transfected using polyethyleneimine (PEI, Polysciences) transfection reagent. Freestyle 293-F cells cell pellets expressing Flag-hEya3 constructs were lysed in 20 mM Tris pH 8, 400 mM NaCl, 0.5 mM TCEP, 1 mM MnCl_2_, 0.1% Triton X-100, and 1X PIC. Lysis was done on ice with sonication using 3 x 10 second pulses. Lysate was centrifuged in an ultracentrifuge at 22,000 RPM and 4 °C for 45 minutes. Supernatant was loaded onto equilibrated α-Flag resin and incubated while rotating at 4 °C for 2 hours. Flag-Eya3 was eluted by incubating the resin with 0.2 mg/mL Flag peptide in elution buffer containing 20 mM pH 8 Tris, 250 mM NaCl, 1 mM MnCl_2_ and 0.5 mM TCEP. Eluates were analyzed with SDS-PAGE and Coomassie stain. Peaks were combined, concentrated, and stored at -80 °C.

Flag-PP2A-B55 was expressed and purified from cells transfected with pCAG-Flag-B55, using the same method as Flag-hEya3. Flag-PP2A-B55 protein also brings down endogenous PP2A-A and PP2A-C subunits from the cell. The eluted PP2A holoenzyme was concentrated and further purified using Superdex S200 Increase 10/300 GL column with buffer (20 mM Hepes pH 8, 150 mM NaCl, 1 mM MnCl_2_ and 0.5 mM TCEP). Proteins were analyzed by SDS-PAGE and fractions containing the PP2A-B55α holoenzyme were pooled, concentrated, aliquoted, and stored at -80 °C.

His-B55 was expressed in Hi5 cells. Cells were lysed in 50 mM pH 8 Tris, 100 mM NaCl, 1X protease inhibitor cocktail (PIC) (Thermo Scientific), PMSF, Leupeptin, Pepstatin A, and 1 mM dithiothreitol (DTT). Cells were sonicated on ice using 6 x 20 second pulses with 60 second off periods. Lysate was centrifuged at 20,000 RPM for 60 minutes in an ultracentrifuge. Equilibrated nickel resin was incubated with cell lysate while rotating for 2 hours at 4 °C. Resin was washed with lysis buffer with 20 mM imidazole and eluted with 500 mM imidazole. PP2A-C and PP2A-A were expressed and purified as previously described (46) from SF9 cells and *E. coli* respectively.

His-B56γ1 was expressed and lysed the same as GST-mEya3. His-B56γ1 was incubated with equilibrated nickel resin at room temperature. The resin was washed with lysis buffer containing 20-300 mM imidazole. The His-B56γ1 was then eluted with 300-500 mM imidazole and analyzed with SDS-PAGE and Coomassie staining. Elution fractions containing His-B56γ1 were concentrated, aliquoted, and stored at -80 °C.

### Cryo-EM structural determination

The PP2A-B55 + Eya3 complex was prepared by incubating 10 μM (final concentration) of purified PP2A-B55 with 15 μM Eya3. The PP2A-B55 + B55i complex was prepared by incubating 14 μM of purified PP2A:B55α with 30 μM B55i peptide. Immediately prior to blotting and vitrification (Vitrobot MK IV, 4 °C, 100% relative humidity, blot time 2-3 s), CHAPSO (3-([3-cholamidopropyl]dimethylammonio)-2-hydroxy-1-propanesulfonate) was added to a final concentration of 0.1% (w/v).

For the apo PP2A-B55 complex, images were collected at Case Western Reserve University cryo-EM core facility using a Titan Krios that was operated at 300 keV and equipped with a K3 direct detector. A total of 4272 movies were recorded, with a super-resolution pixel size of 0.535 Å, a defocus range of −1∼−2.5 μm, and 50 frames per movie with a total dose of ∼54 electrons/Å^2^. Patch motion correction (2x binned) and patch CTF estimation were performed in cryoSPARC v4.4.0. A total of ∼2.4 million particles were automatically picked using the Blob Picker. Those particles were extracted with a box size of 216 × 216 pixels and subjected to several rounds of 2D classification. ∼884,147 particles were selected for ab initio reconstruction and several runs of heterogeneous refinement. From this, 233,311 good particles were chosen for homogenesis refinement and nonuniform refinement. The final map reached a resolution of ∼3.16 Å, as determined by the gold-standard Fourier shell correlation (FSC) at a cutoff of 0.143.

For the PP2A-B55 + Eya3 and PP2A-B55 + B55i complexes, images were collected at NCEF using a 300 keV Titan Krios microscope, equipped with a K3 direct detector (Gatan). A total of 5520 (for Eya3) or 5462 (for B55i) movies were recorded with a pixel size of 0.855 Å, a defocus range of −0.6∼−2.0 μm, and 50 frames per movie, with a total dose of ∼50.02 electrons/Å^2^. Patch motion correction and patch CTF estimation were done in cryoSPARC v4.4.0. For the eya3 dataset, a total of ∼2.5 million particles were automatically picked using the Blob Picker. Those particles were extracted with a box size of 320 × 320 pixels and subjected to several rounds of 2D classification. ∼707,280 particles were selected for ab initio reconstruction and several runs of heterogeneous refinement. A subset of 159,895 good particles was then selected for homogenesis refinement and nonuniform refinement, yielding a map with a resolution of ∼3.71 Å, as determined by the gold-standard Fourier shell correlation (FSC) at a cutoff of 0.143. For the B55i dataset, a total of ∼1.8 million particles were automatically picked using the Blob Picker and extracted with a box size of 320 × 320 pixels. Those particles were used for the ab initio reconstruction of 5 initial models. These initial models were heterogeneously refined, and the best group with 171,051 particles was selected for further homogeneous refinement and nonuniform refinement. The final map reached a resolution of ∼3.5 Å, based on the gold-standard Fourier shell correlation (FSC) at a cutoff of 0.143.

All models were built in Coot. To aid subunit assignment and model building, we took advantage of the reported PP2A-B55 + FAM122A (PDB code: 8SO0) which was fitted into the PP2A-B55, PP2A-B55 + Eya3 and PP2A-B55 + B55i density map with appropriate main chain and side chain adjustment. The final rounds of model refinement were carried out by real-space refinement in PHENIX.

### Microscale Thermophoresis (MST)

His-B55 or His-B56γ1 labeled with Red-His-NTA (NanoTemper) was added to serial dilutions of mEya3 in PBS with 0.05% Tween 20. The samples were then read with a Monolith NT.115^Pico^ (NanoTemper) according to the manufacturer’s instructions.

### Fluorescence Polarization

His-B55 was serially diluted in 50 mM pH 8 Tris, 100 mM NaCl, and 1 mM DTT. His-B55 was incubated with 3 nM of FITC-B55i peptide (GL Biochem) at room temperature on a horizontal shaker for 10 minutes prior to reading. Fluorescence polarization was measured using the EnVision HTS Plate Reader (PerkinElmer).

### Malachite Green Phosphatase Assay

100 μM pT58-P59A phosphosubstrate was incubated for 45 minutes at 37 °C with PP2A-B55 in 30 mM HEPES pH 7.0, 150 mM NaCl, and 1 mM MnCl_2_. Phosphatase reactions were then quenched with 0.01% tween-20, 0.034% malachite green, 10 mM ammonium molybdate, 1 N HCl, and 3.4% ethanol. Final reaction solutions were analyzed with a Molecular Devices plate reader at 630 nm.

### 3-O-methylfluorescein phosphate (OMFP) Assay

25 nM of PP2A-B55 holoenzyme was with or without 1 μM Eya3 NTD in 30 mM HEPES pH 7.0, 150 mM NaCl, 1 mM MnCl2, 1 mM DTT, 0.01% triton X-100, 0.1 mg/ml BSA was incubated at room temperature for ten minutes to bind. The reaction was carried out at 37 °C for 30 minutes. Fluorescent OMFP, corresponding to phosphatase activity, was read using a BioTek Synergy H1 plate reader (excitation at 485 nm and emission at 515 nm).

### Western Blot

Western blot was performed as previously described (46). Primary antibodies used include: α-Myc (rabbit, Abcam Y69), α-Myc pT58 (rabbit, ABM Y011034), and α-GAPDH (mouse, GeneTex GT239). These antibodies were validated as previously described (46).

### Mass Photometry (MP)

hEya3 and PP2A-B55 were combined in 20 mM pH 8 Tris, 250 mM NaCl, 1 mM MnCl_2_ and 0.5 mM TCEP at a molar ratio of 1.5:1. The protein complex was incubated on ice in the presence or absence of 25 μM B55i peptide (GL Biochem). Immediately prior to reading, the solution was diluted to 100 nM. Samples were analyzed on the Refeyn TwoMP (Refeyn).

## Data availability

Cryo-EM structures of the PP2A-B55 apo, + Eya3, + B55i were deposited in the Protein Data Bank with ID 9MZW, 9N0Y, and 9N0Z, respectively. The corresponding EM maps were deposited in the EMDB with ID EMD-48770, EMD-48798, and EMD-48799, respectively.

## Supporting Information

This article contains supporting information.

## Acknowledgements

We thank Drs. Rebecca Page and Wolfgang Peti for helpful discussions. We thank the University of Colorado Anschutz Medical Campus Cryo-EM Core Facility, the Biophysics Core Facilities, and the University of Colorado Boulder Cryo-EM Core Facility, and the core facility staff members for help with data acquisition.

## Funding and Additional Information

Research reported was supported by the National Institute of Health under award number R01CA221282 (H.L.F. and R.Z.), R35GM145289 (R.Z.), R01CA224867 (H.L.F.), R01CA275187 (H.L.F.), R01 CA240993 (D.J.T.), R01GM154832 (D.J.T.), RM1GM142002 (D.J.T.). It was also supported by Cancer League of Colorado grant (R.Z.) and Jordan’s Guardian Angels (D.J.T.). C.A. and L.W. were supported by the NIH NRSA T32CA174648 and T32CA190216, respectively. The Anschutz Medical Campus Cryo-EM Core Facility was supported in part by NIH P30CA046934.

## Supplementary Figure Legends

**Supplementary Figure 1. Eya3 and His-B55 purification.** Eya3 purified from *E. coli* (A) and His-B55 purified from insect cell (B) are shown on Coomassie stained gel.

**Supplementary Figure 2. Structural determination of the PP2A-B55 + Eya3 complex.** A) PP2A-B55 complex purified from FreeStyle HEK293 cells shown on Coomassie stained gel. B) Flag-tagged human Eya3 purified from FreeStyle HEK293 cells shown on Coomassie stained gel. C) A representative negative stain image of the PP2A-B55 + Eya3 complex. D) A representative cryo image of the PP2A-B55 + Eya3 complex. E) 2D classification of the cryo-EM data.

**Supplementary Figure 3.** A) Workflow of the structural determination process. B) Golden Standard FSC curves show the overall resolution of the structure. C) Cryo-EM map of the structure colored by local resolutions. A ribbon diagram of the structure is also shown to illustrate the orientation of the local resolution map. D) Density for residues 73-84 of Eya3.

**Supplementary Figure 4. Structural determination of the PP2A-B55 + B55i complex.** A) A representative negative stain image of the PP2A-B55 + B55i complex. B) A representative cryo image of the PP2A-B55 + B55i complex. C) 2D classification of the cryo-EM data. D) Workflow of the structural determination process. E) Golden Standard FSC curves show the overall resolution of the structure. F) Cryo-EM map of the structure colored by local resolutions. A ribbon diagram of the structure is also shown to illustrate the orientation of the local resolution map. G) Density for the C-terminal helix (residues 22-35) of B55i.

**Supplementary Figure 5. Structural determination of the apo PP2A-B55 complex.** A) A representative negative stain image of the apo PP2A-B55 complex. B) A representative cryo image of the apo PP2A-B55 complex. C) 2D classification of the cryo-EM data. D) Workflow of the structural determination process. E) Golden Standard FSC curves show the overall resolution of the structure. F) Cryo-EM map of the structure colored by local resolutions. A ribbon diagram of the structure is also shown to illustrate the orientation of the local resolution map. G) Density for the C-terminal tail in the PP2A-B55 + Eya3 and apo structures.

**Supplementary Table 1.** Data statistics for structural determination and refinement.

