## Supplementary Figures for "Cryo-EM structures reveal the PP2A-B55α and Eya3 interaction that can be disrupted by a peptide inhibitor"

Supplementary Figure 1

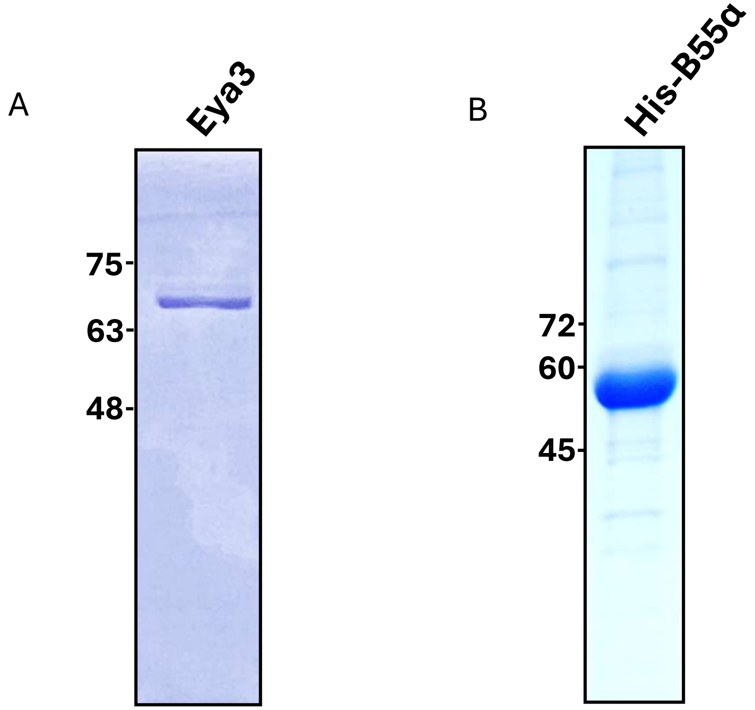

Supplementary Figure 2

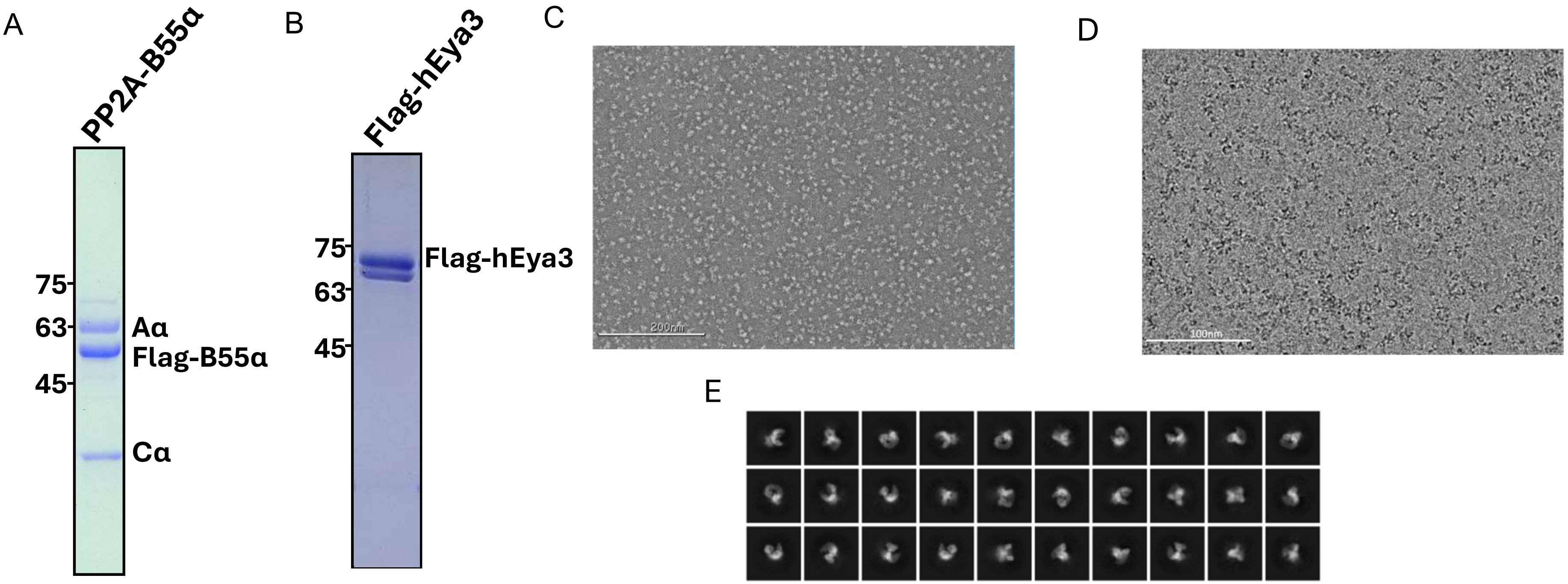

#### Supplementary Figure 3

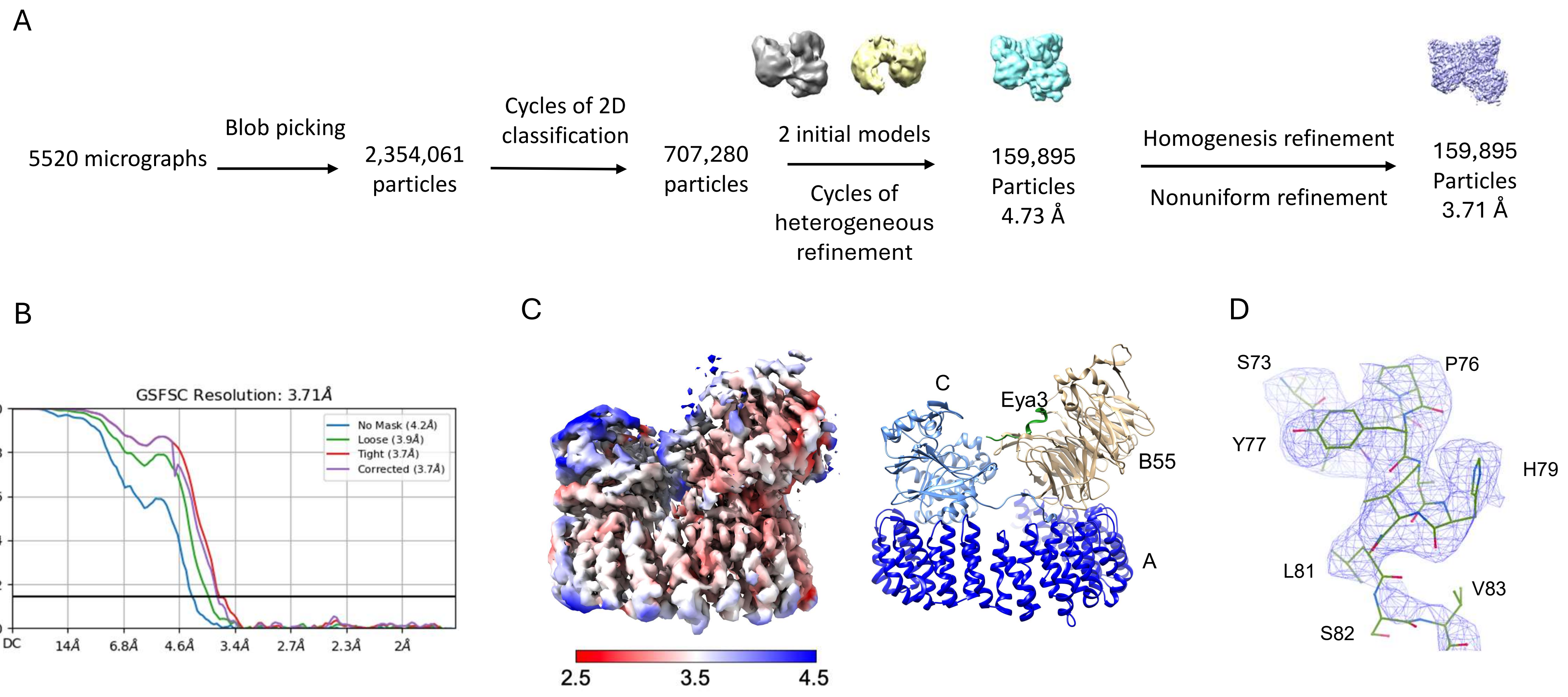

Supplementary Figure 4

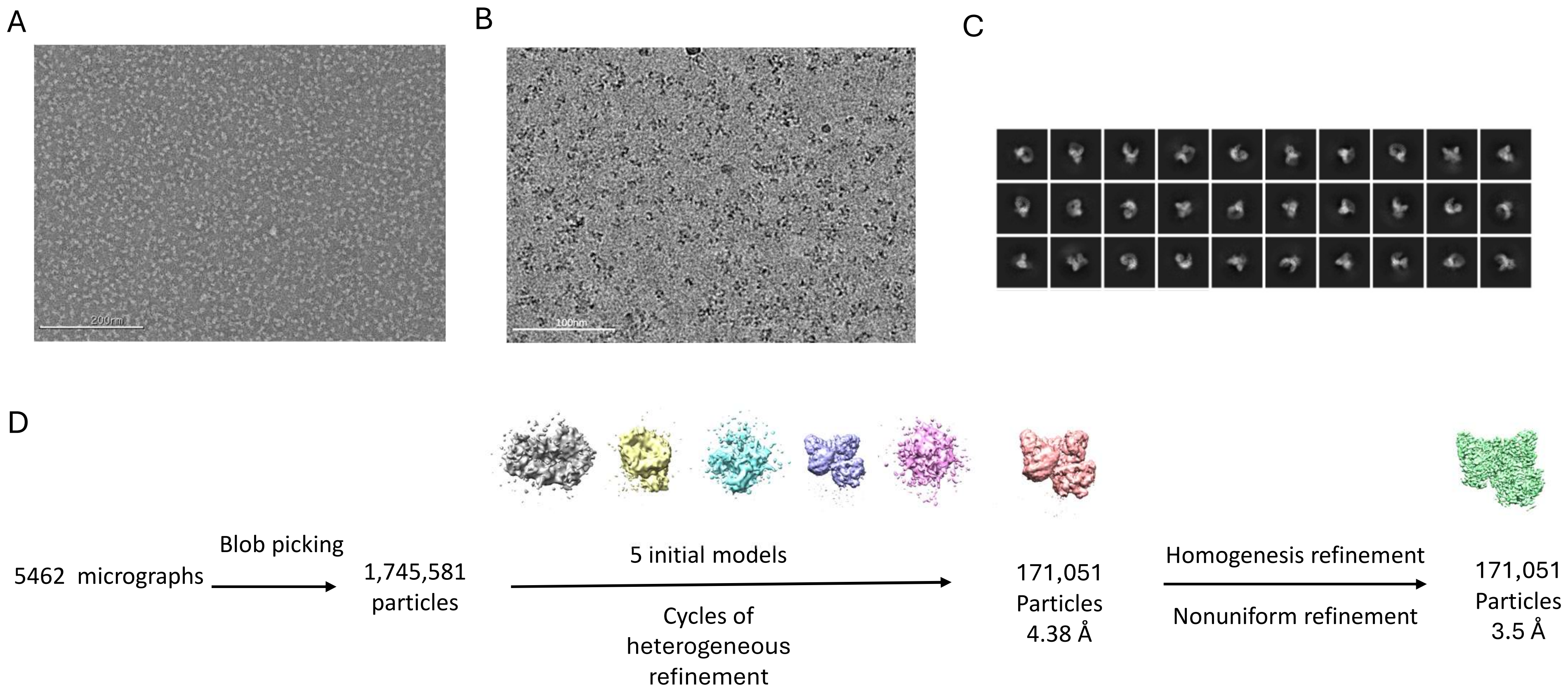

Supplementary Figure 4

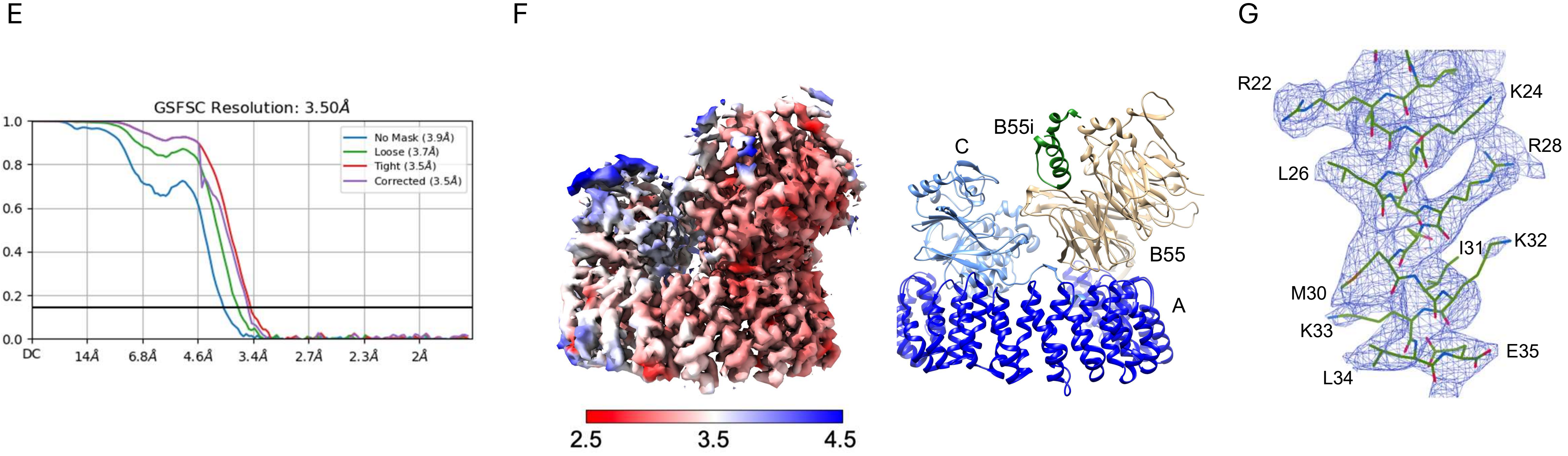

Supplementary Figure 5

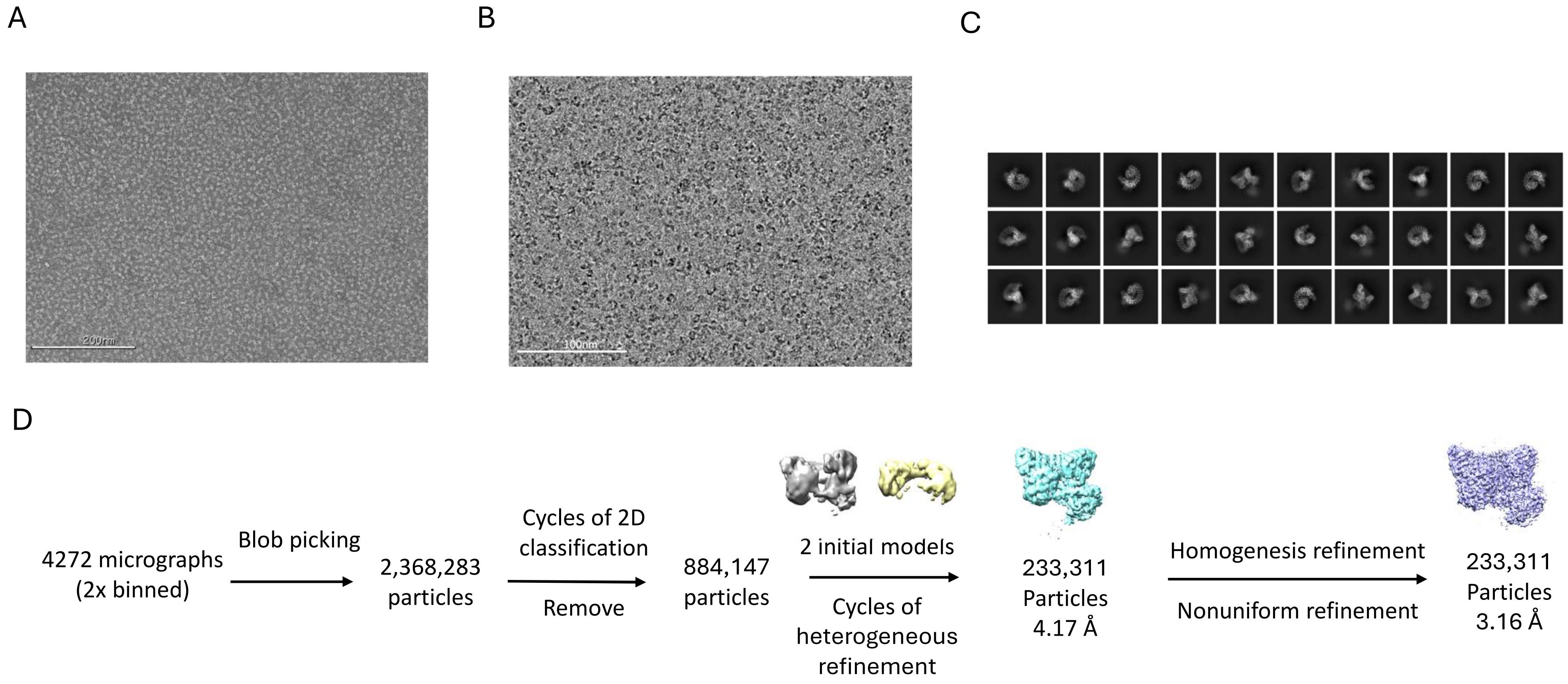

### Supplementary Figure 5

E

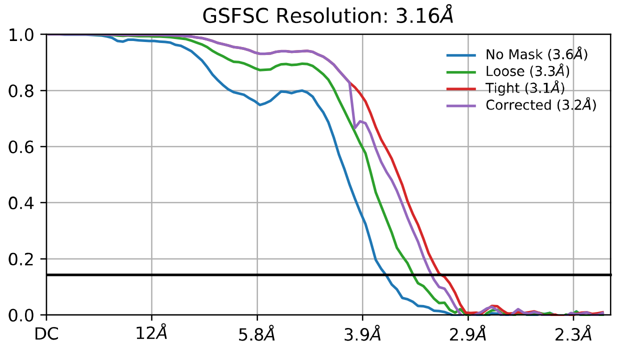

F

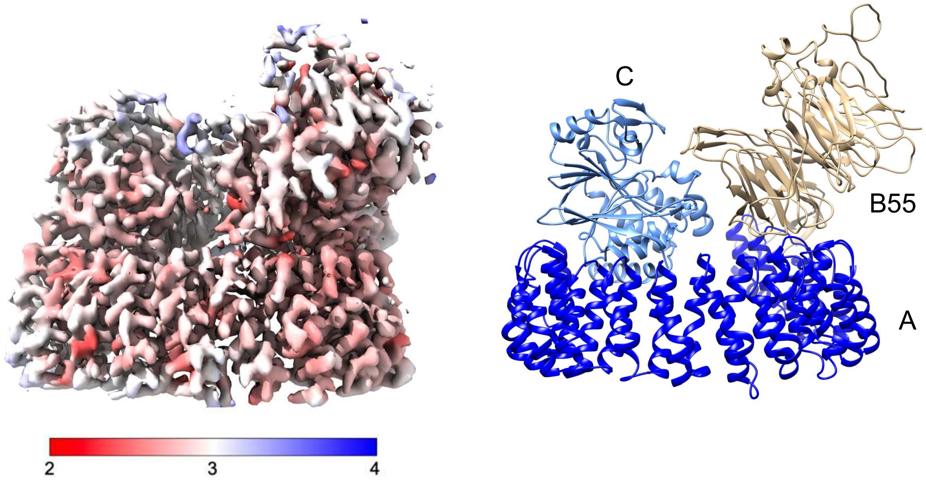

G

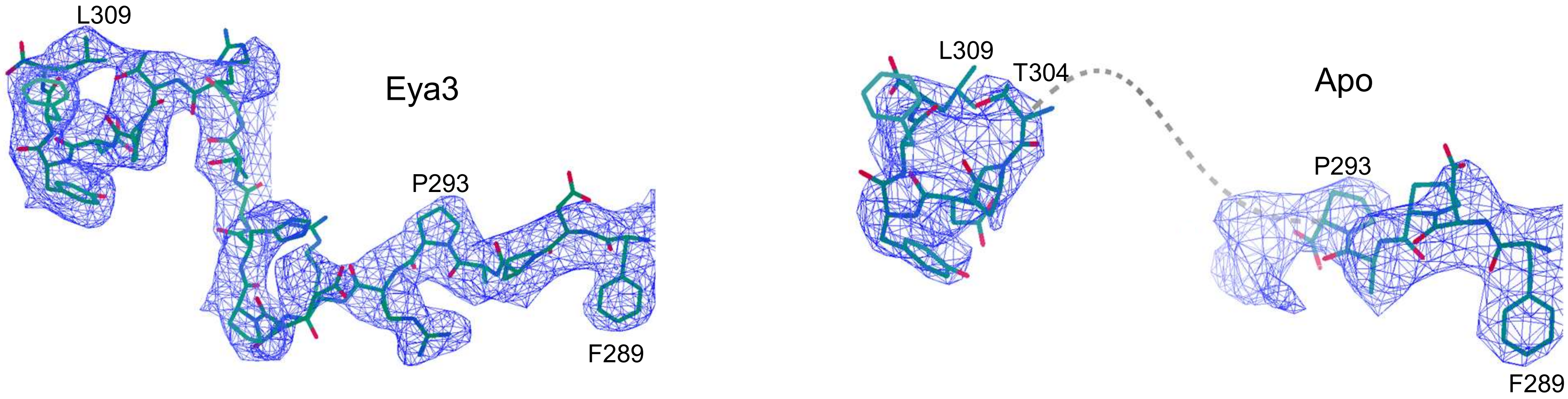

### Supplementary Table 1

| | PP2A-B55a (EMD-48770) (PDB 9MZW) | PP2A-B55 + Eya3 (EMD-48798) (PDB 9N0Y) | PP2A-B55 + B55i (EMD-48799) (PDB 9N0Z) |
|---|---|---|---|
| **Data collection, processing** | | | |
| Magnification | 81,000 | 105,000 | 105,000 |
| Voltage (Kv) | 300 | 300 | 300 |
| Exposure dose ((e-/Å2)) | 54 | 50.02 | 50.02 |
| Defocus range (µm) | −1~−2.5 | −0.6 ~ −2.0 | −0.6 ~ −2.0 |
| Pixel size (Å) | 0.535 | 0.855 | 0.855 |
| Movies collected | 4272 | 5520 | 546200 |
| Symmetry imposed | C1 | C1 | C1 |
| Initial particle images (no.) | 2,368,283 | 2,354,061 | 1,745,581 |
| Final particle images (no.) | 233,311 | 159,895 | 171,051 |
| Map resolution (Å) | 3.16 | 3.71 | 3.5 |
| FSC threshold | 0.143 | 0.143 | 0.143 |
| Map resolution range | 2.3-4.2 | 2.5-5.5 | 2.5-4.9 |
| **Refinement** | | | |
| Initial model used | 8SO0 | 8SO0 | 8SO0 |
| model composition | | | |
| Chains | 3 | 4 | 4 |
| Atoms | 10479 | 10723 | 13248 |
| Residues | 1312 | 1342 | 1355 |
| R.m.s. deviations | | | |
| Bonds (Å) | 0.004 | 0.004 | 0.005 |
| Bond angles (º) | 0.627 | 0.693 | 0.674 |
| **Validation** | | | |
| MolProbity score | 1.94 | 2.14 | 2.44 |
| Clash score | 13.72 | 12.13 | 26.63 |
| Rotamer outliers (%) | 0.00 | 0.08 | 0.17 |
| Ramachandran plot | | | |
| Favored (%) | 95.71 | 90.18 | 90.85 |
| Allowed (%) | 4.29 | 9.82 | 9.15 |
| Disallowed (%) | 0.00 | 0.00 | 0.00 |
